# Does the Ice Age legacy end in Central Europe? The shrinking distributions of glacial relict crustaceans

**DOI:** 10.1101/2022.11.23.517644

**Authors:** Kęstutis Arbačiauskas, Carl Smith, Asta Audzijonyte

**Affiliations:** Nature Research Centre, Vilnius, Lithuania; Department of Zoology, Life Science Center, Vilnius University, Vilnius, Lithuania; Department of Ecology and Vertebrate Zoology, University of Łódź, Łódź, Poland; Institute of Vertebrate Biology of the Czech Academy of Sciences, Brno, Czech Republic

**Keywords:** biological invasions, climate change, eutrophication, glacial relicts, *Mysis relicta*, *Monoporeia affinis*, *Pallaseopsis quadrispinosa*, Ponto-Caspian crustaceans

## Abstract

1. Glacial relict mysid and amphipod crustaceans are characterised by their affinity for cold and well-oxygenated waters and inability to disperse upstream or by external agents. These crustaceans occur in large and deep lakes of Northern and Central Europe and North America, their distributions shaped by glaciation events. In Europe, along the southern edge of their distribution (Germany, Poland, Lithuania, Belarus), glacial relict crustaceans are threatened by eutrophication and global warming.
2. This study assesses the status of three glacial relict malacostracan species in Lithuania; the amphipods *Monoporeia affinis* and *Pallaseopsis quadrispinosa*, and mysid *Mysis relicta*, and models their abundance as a function of environmental variables and the presence of invasive Ponto-Caspian mysids and amphipods.
3. Our results revealed that *M. affinis* is likely extinct in the country, whereas *M. relicta* was found in only 9 out of 16 locations from which it was previously recorded. The distribution of *P. quadrispinosa* appears to be shrinking.
4. Lake depth and water flowthrough intensity were significantly and positively associated with the relative abundance of relict mysids and amphipods, but no association was found with lake size or the presence of invasive Ponto-Caspian crustaceans.
5. We conclude that urgent action to mitigate the effects of nutrient run-off is needed to improve the status of glacial relict and other species that require good water quality. We also propose the re-introduction of glacial relict species in Lake Drūkšiai, where they went extinct during the operation of the Ignalina nuclear power plant that heated the lake, but where deep-water environmental conditions have improved following the powerplant closure in 2010.

## 1. Introduction

Several species of malacostracan crustaceans, mysids and amphipods in particular, have colonised the inland waters of northern Europe and North America through waterways created by post-glaciation events (Segerstråle, 1957; Holmquist, 1966). Because these species cannot disperse by external agents, their occurrences, confined to the formerly glaciated areas, contributed to the formulation of the idea that past glaciation events might have shaped current species and population distributions (Lovén, 1862; Högbom, 1916). These crustacean species are characterised by a requirement for cold and well-oxygenated water and are often referred to as glacial relicts (Lovén, 1862; Thienemann, 1925; Audzijonyte & Väinölä, 2005). They typically inhabit large, oligotrophic and mesotrophic lakes of North America and Europe. In Europe, glacial relict species are common in Scandinavia, with their southern distribution limit extending to Germany, Poland, Lithuania and Belarus (Gasiūnas, 1959; Köhn & Waterstraat, 1990; Żmudziński, 1990).

In suitable habitats, glacial relict crustaceans can be found in high abundances and play a significant ecological role in pelagic food webs, often comprising most of the diet of commercially valuable inland fish species, such as vendace, whitefish and others (Lasenby, Northcote & Fürst, 1986; Sandlund, Næsje & Jonsson, 1992; Scharf et al., 2008). However, during the 20^th^ century and especially during the last few decades, glacial relict crustaceans in the southern part of their distribution have faced increasing anthropogenic pressure, with extinctions reported in many locations. For example, the most sensitive glacial relict species *Monoporeia affinis* (Lindström, 1855), is now considered extinct in Germany (Köhn & Waterstraat, 1990) and Poland (Żmudziński, 1995) but is still known to occur in one lake in Belarus (Vezhnavets V.V., 2021, pers. comm.). The other two glacial relict species found are a mysid, *Mysis relicta* Loven, 1862 (sensu stricto, see Audzijonyte & Väinölä, 2005), and amphipod, *Pallaseopsis quadrispinosa* (G.O. Sars, 1867). These species are often listed in national Red Data books in Central Europe.

Two main factors responsible for the decrease in abundance of glacial relict crustaceans are the eutrophication of lakes and global warming, which reduce the availability of cool oxygen-rich habitats typical of the deeper layers of large lakes. Another potential factor is the introduction or invasions of mysids and amphipods from the Ponto-Caspian region. These Ponto-Caspian malacostracans are among the major invaders of European inland ecosystems (JaŻdŻewski, 1980; Bij de Vaate et al., 2002) and are known to be highly successful competitors and predators (Dick & Platvoet, 2000; Arbačiauskas & Gumuliauskaitė, 2007). Despite recognition of their negative ecological impacts, the role of the Ponto-Caspian crustaceans in the disappearance of glacial relict species has not been formally tested. In this study, data were compiled from historical records and recent investigations of the distribution and abundance of glacial relict and Ponto-Caspian malacostracan species in 26 Lithuanian lakes and used to map the current distribution of glacial relict species and model the predictors of their abundance.

## 2. Methods

### 2.1. Species

Original investigations of glacial relict species distributions in Lithuania were conducted in the mid-20^th^ century and were based on extensive sampling of all lakes potentially suitable for these species (Gasiūnas 1959; Grigelis 1980) (Appendix). The three glacial relict malacostracan species recorded in Lithuania and studied here are the relict mysid *Mysis relicta* and the amphipods *Monoporeia affinis* and *Pallaseopsis quadrispinosa*. Historically, *M. affinis* and *M. relicta* have been recorded in 2 and 16 lakes across Lithuania, respectively, while *P. quadrispinosa* was detected in 46 lakes (Appendix). The introduced non-indigenous Ponto-Caspian crustacean species considered in this study include the mysid *Paramysis lacustris* (Czerniavsky, 1882) and amphipods *Pontogammarus robustoides* (G.O. Sars, 1894), *Obesogammarus crassus* (G.O. Sars, 1894) and *Chaetogammarus warpachowskyi* (G.O. Sars, 1894). These species were first introduced into Lithuanian waters between 1960-1961 and are currently known in over 30 lakes (Arbačiauskas et al., 2011; Arbačiauskas et al., 2017; this study).

### 2.2. Sampling, occurrence and relative abundance of malacostracan species

To assess recent changes in the occurrence of glacial relict species, 26 out of 46 lakes previously inhabited by glacial relicts were sampled during 1998-2021 (Table 1, Figure 1). This survey included all lakes known to have been inhabited by the mysid *M. relicta* and the amphipod *M. affinis*. Sampling for glacial relicts and Ponto-Caspian malacostracans was performed in depths between 5 to 30 meters using a modified 70 cm wide epibenthic dredge. The dredge was slowly pulled from a boat for 20-30 meters, targeting the deepest and shallower parts of the lake in separate trawling events. If *M. relicta* and *M. affinis* were not known to occur in the lake and sampling was focused on *P. quadrispinosa* only, species occurrence was first assessed using a hand net in wadable depths late in the autumn season when water temperatures were low, and the species is known to occur at shallower depths. If this sampling approach did not yield results, trawling was conducted. Data on the presence of Ponto-Caspian crustaceans in lakes was based on the same sampling surveys and records from earlier studies (Arbačiauskas et al., 2011; Arbačiauskas et al., 2017).

**Table 1.** Occurrence and relative abundance of glacial relict crustaceans in Lithuanian lakes investigated during 1998-2021. Relict species *Mysis relicta* (M.r.), *Monoporeia affinis* (M.a.) and *Pallaseopsis quadrispinosa* (P.q.), year of sampling and relative abundance in brackets are provided, abundance classes: absent (0), present (1), moderately abundant (2), abundant (3). Ponto-Caspian species (P.C.s.) occurring in a lake: *Paramysis lacustris* (P.l.) *Pontogammarus robustoides* (P.r.), *Obesogammarus crassus* (O.c.) and *Chaetogammarus warpachowskyi* (C.w.).

| No | Lake | M.r. | M.a. | P.r. | P.C.s. |
| --- | --- | --- | --- | --- | --- |
| 1 | Aisetas | 1999 (3), 2003 (3), 2021 (3) |  | 1999 (2), 2003 (2), 2021 (2) |  |
| 2 | Akmena |  |  | 1998 (0) |  |
| 3 | Asalnai | 2003 (2) |  | 2003 (2) | P.l., P.r. |
| 4 | Asveja | 1999 (3), 2003 (3), 2021 (3) |  | 1999 (3), 2003 (3), 2021 (3) | P.r., O.c. |
| 5 | Baltieji Lakajai | 1999 (0) |  | 1999 (3), 2003 (2), |  |
| 6 | Baluošai | 2003 (3), 2021 (3) |  | 2003 (2), 2021 (2) |  |
| 7 | Baluošas | 1999 (3), 2003 (1), 2021 (1) |  | 1999 (3), 2003 (3), 2021 (1) | P.l. |
| 8 | Čičirys |  |  | 2003 (1) |  |
| 9 | Drūkšiai | 2003 (0) |  | 2003 (0) | P.l. |
| 10 | Dusia | 1999 (1), 2000 (1), 2002 (1), 2005 (0) | 1999 (0), 2000 (0) | 1999 (3), 2000 (2), 2002 (1), 2005 (0) | P.l., P.r., O.c., C.w. |
| 11 | Galstas |  |  | 2014 (2) | P.r. |
| 12 | Galvė |  |  | 2020 (1) |  |
| 13 | Ilgai |  | 1999 (0) | 1999 (3) |  |
| 14 | Luodis |  |  | 1995 (1), 2008 (0) |  |
| 15 | Luokesai | 2003 (0) |  | 2003 (2) |  |
| 16 | Lūšiai | 2003 (3), 2013 (3), 2021 (2) |  | 2003 (3), 2013 (3), 2021 (3) | P.l., P.r., C.w. |
| 17 | Peršokšnai | 2003 (2), 2021 (0) |  | 2003 (1), 2021 (0) |  |
| 18 | Siesartis | 2003 (0) |  | 2003 (0) |  |
| 19 | Šakarvai | 1998 (3), 2003 (3) |  | 1998 (3), 2003 (3) | P.l., P.r., C.w. |
| 20 | Šventas |  |  | 2018 (0) |  |
| 21 | Tauragnas |  |  | 1998 (3), 2003 (3) |  |
| 22 | Ūkojas | 1999 (3), 2003 (0) |  | 1999 (2), 2003 (1) |  |
| 23 | Vencavas |  |  | 1999 (3) |  |
| 24 | Vištytis |  |  | 2014 (2) | P.r. |
| 25 | Zarasas | 2003 (3), 2021 (3) |  | 2003 (3), 2021 (3) |  |
| 26 | Žeimenys | 2003 (3) |  | 2003 (2) | P.l., P.r., C.w. |

**Figure 1.**
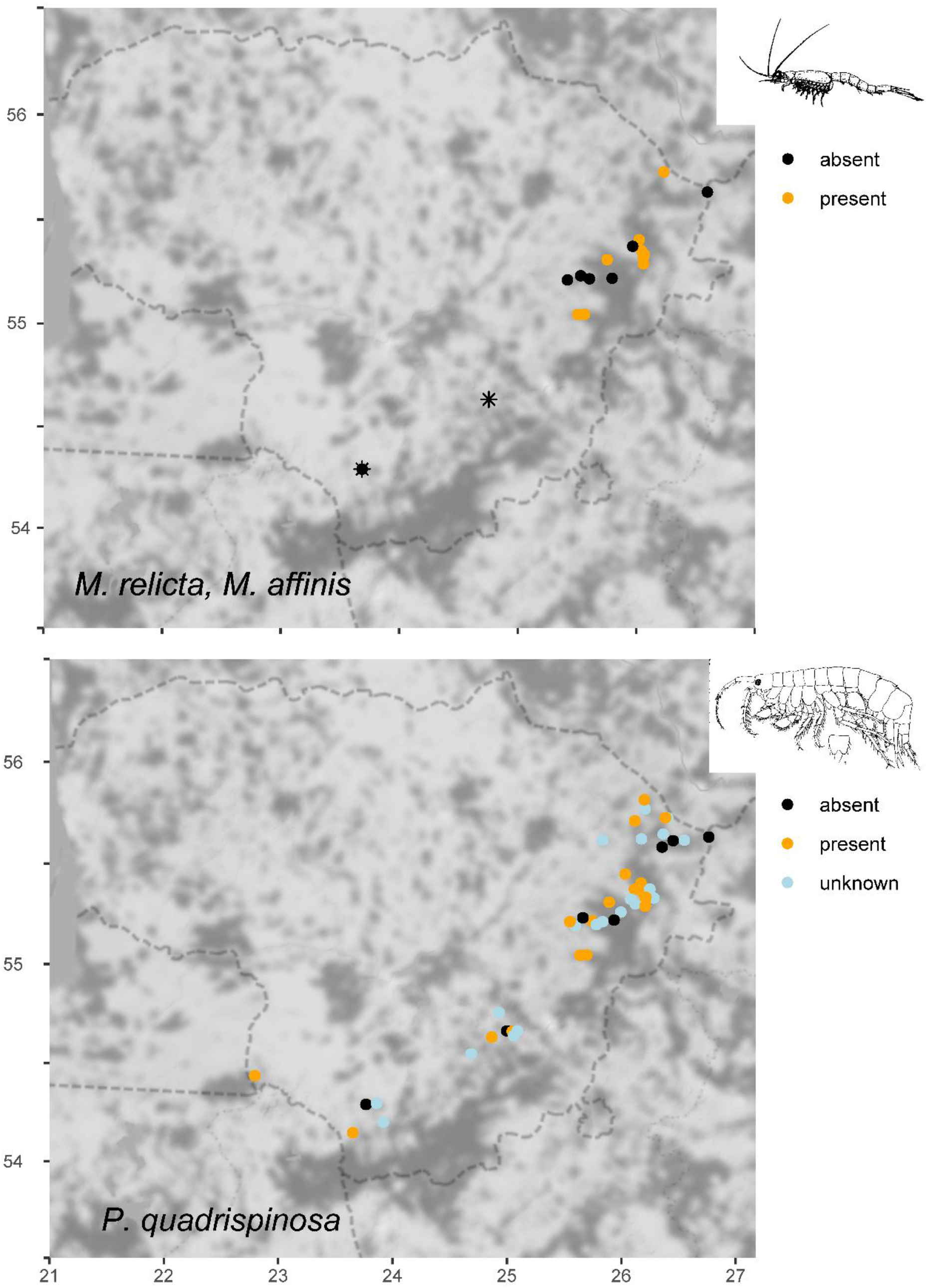
Location of Lithuanian lakes with historical records of glacial relict crustaceans indicating the recent occurrence of these crustaceans. The amphipod *Monoporeia affinis* (asterisks) and the mysid *Mysis relicta* (circles) occurred in 2 and 16 lakes, respectively. Recently, *M. affinis* became extinct from both localities and *M. relicta* survived in 9 localities (above). The amphipod *Pallaseopsis quadrispinosa* occurred in 46 lakes. Recently it was not recorded in 7 lakes, was found to be present in 19 lakes, while 20 localities remained unchecked (below).

The relative abundance of glacial relict crustaceans in this study was assessed based on the number of individuals sampled in one trawling event. Three relative abundance categories were used: *abundant*, when the number of individuals found in at least one trawling event at the site was >100; *moderately abundant*, with 11 to 100 individuals per trawling event; and *rare*, when no one trawling event yielded more than ten individuals. On a few occasions, when the presence of *P. quadrispinosa* was detected by hand net sampling, the presence of ≤5 or >5 specimens per 5 min sampling effort was interpreted as indicating a moderate (≤5) or abundant (>5) population.

### 2.3. Statistical analysis

Five environmental variables were examined as predictors of glacial relict crustacean abundance: lake surface area, maximum lake depth, average lake depth, lake flowthrough and the presence of Ponto-Caspian crustaceans. Lake flowthrough measures the proportion of water replaced annually by inflowing and outflowing rivers and is an important contributor to the oxygenation of deeper water layers.

Consequently, flowthrough was used as a proxy for oxygen conditions in a lake, given that data on oxygen levels in deeper water layers were unavailable. Environmental variables about lake depth, area and flowthrough were obtained from the data archive of the Institute of Geology and Geography (Geologijos ir geografijos institutas, 2002). The full dataset used in the analysis is available in Table 1 and Appendix.

To assess which of the variables predicted the relative abundance of glacial relict crustaceans, Generalized Linear Mixed Models (GLMM), as implemented in libraries *nlme* (Pinheiro et al., 2020) and *glmmTMB* (Brooks et al., 2017) in the R environment (version 4.2.1; R Core Team, 2022), were fitted. Before model fitting, a data exploration was undertaken following the protocol described in Ieno & Zuur (2015). The data were examined for outliers in the response and explanatory variables, homogeneity and zero inflation in the response variable, collinearity between explanatory variables and the nature of relationships between the response and explanatory variables. Sample variograms failed to show spatial dependency in the data, and an autocorrelation function plot failed to show temporal autocorrelation. Separate analyses were performed for *M. relicta* and *P. quadrispinosa* relative abundance. Because sampling among years was highly uneven, year was treated as a category with two levels: samples from the 20^th^ century (Period 1) or 21^st^ century (Period 2). Maximum and average lake depths were highly correlated, so only average depth was included in models.

The relative abundance of relict crustaceans was modelled using a Generalized Linear Mixed Model (GLMM) with Conway-Maxwell Poisson distribution; this distribution, rather than a Poisson, was employed to accommodate underdispersion in the response variable. The initial full model was defined as:

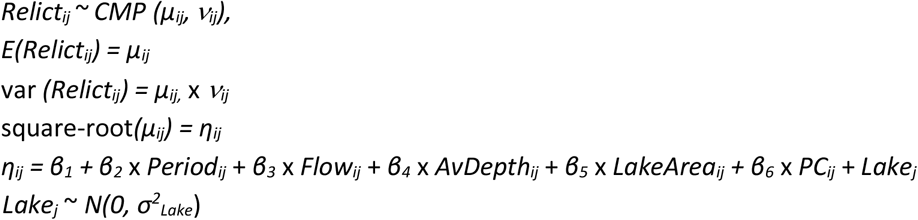

where *Relict*_*ij*_ is the relative abundance of relict crustaceans (amphipod or mysid) in sample *i* from lake *j*, which was assumed to follow a Conway-Maxwell-Poisson distribution with an expected abundance (*E*) in each sample with mean *μ*_*ij*_ and variance *μ*_*ij*,_ x *v*_*ij*_ (where *v* is a dispersion parameter) and a square-root link function. The fixed effects were the period of sample collection (*Period*_*ij*_), lake flowthrough (*Flow*_*ij*_), average lake depth (*AvDepth*_*ij*_), lake area (*LakeArea*_*ij*_) and presence of Ponto-Caspian crustaceans (mysids or amphipods) in the lake (*PC*_*i,j*_) at the time of sampling, while *β*_*1*_ to *β*_*6*_ are model coefficients to be estimated. To accommodate the fact that lakes were sampled repeatedly and the abundance of relict crustaceans differed among lakes, the random intercept *Lake*_*j*_ was included in models to introduce a correlation structure between observations for different samples from the same lake, with variance σ^2^_*lake*_ distributed normally and with a mean of zero (categorical, 26 levels). We could not perform a time series or survival analysis on this dataset due to a limited number of sampling years per lake. The optimal fixed structure of a model for each dataset was identified with a backward selection procedure using AIC (Δ AIC ≤ 2) (Akaike, 1973). All datasets and scripts used in these analyses are available on the GitHub repository https://github.com/astaaudzi/relictCrustaceans.

## 3. Results

Studies conducted in the 1950s reported the presence of glacial relict malacostracans in 46 Lithuanian lakes (Figure 1, Appendix). More recent sampling performed in 26 of these lakes, including all lakes with earlier records of *M. affinis* and *M. relicta*, revealed that *M. affinis* is likely to be extinct in Lithuania, as it has not been found in either of the two lakes in which it was earlier known to occur (Dusia and Ilgai, #16 and #20 in the Appendix, asterisks in Figure 1). In the mid-20^th^ century, *M. relicta* was recorded in 16 lakes, whereas recent data suggest that the species now occurs only in 9 lakes (Table 1, Figure 1). The amphipod *P. quadrispinosa* was known to occur in all 26 studied lakes, but sampling failed to detect this species in 7 of the studied lakes. Ten out of 26 investigated lakes currently contain invasive Ponto-Caspian crustaceans, and in two of these lakes, extinctions of glacial relicts were observed (Table 1, Figure 1).

Backward model selection showed that for both relict mysid and amphipod datasets, the best-fitting model included three environmental variables: average depth, flowthrough and time period (20^th^ or 21^st^ century).

Removing the presence of Ponto-Caspian crustaceans from models improved model fit (AIC of 331 versus 180 for amphipods, and AIC of 167 versus 126 for mysids), showing that the presence of Ponto-Caspian crustaceans did not explain the abundance of glacial relicts.

The final models showed relict crustacean relative abundance to be positively associated with lake flowthrough (for amphipods P = 0.009; for mysids P = 0.028) and average lake depth (amphipods and mysids P <0.001) (Table 2, Figures 2, 3). For each species, there was also a significant association with time period, and a reduction in relative abundance associated with the 21^st^ century compared to the 20^th^ century (amphipods P = 0.007; mysids P = 0.004). Fixed effects explained about 20% of the variance in relict crustacean relative abundance in both models (Table 2). In contrast, a lake effect, included as a random term in the models, explained approximately 8-10% of the variance in both models (Table 2: the difference between conditional R^2^, defined by fixed effects, and marginal R^2^, defined by both fixed and random effects).

**Table 2.**
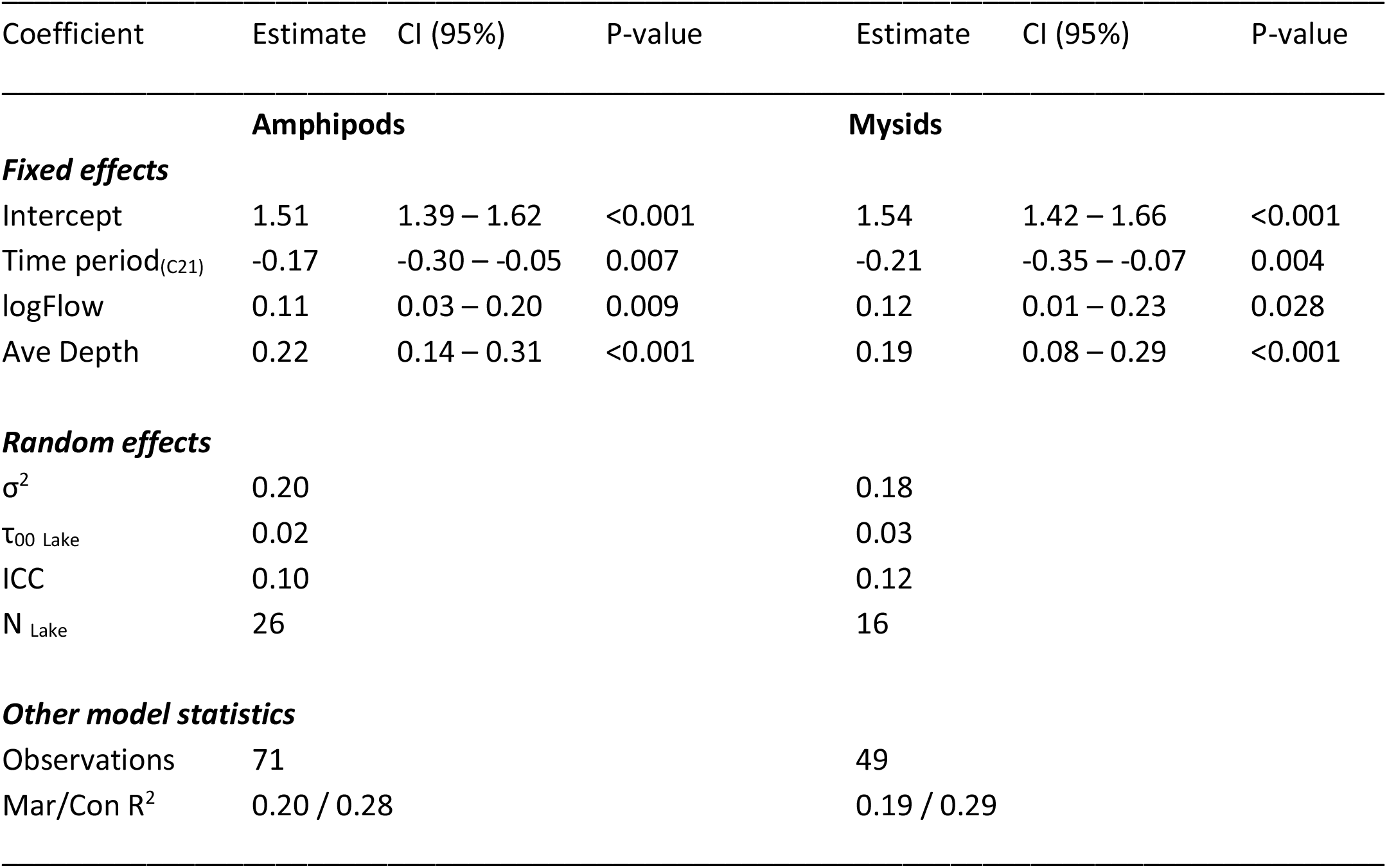
Parameters of the final Conway-Maxwell Poisson generalized linear models for relict mysid *Mysis relicta* and relict amphipod *Pallaseopsis quadrispinosa* relative abundance analyses. Mar/Con R^2^ – marginal and conditional R^2^ values; *σ*^2^ is the mean random effect variance for each model; τ_00 Lake_ is the model between-subject variance, indicating how much different levels of the random term ‘Lake’ differ from each other; ICC is the intra-class correlation coefficient, which is a measure of the degree of correlation within groups; N indicates the number of levels in the random effect ‘Lake’.

| Coefficient | Estimate | CI (95%) | P-value | Estimate | CI (95%) | P-value |
| --- | --- | --- | --- | --- | --- | --- |
| **Amphipods** |  |  |  | **Mysids** |  |  |
| **Fixed effects** |  |  |  |  |  |  |
| Intercept | 1.51 | 1.39 – 1.62 | <0.001 | 1.54 | 1.42 – 1.66 | <0.001 |
| Time period<sub>(C21)</sub> | -0.17 | -0.30 – -0.05 | 0.007 | -0.21 | -0.35 – -0.07 | 0.004 |
| logFlow | 0.11 | 0.03 – 0.20 | 0.009 | 0.12 | 0.01 – 0.23 | 0.028 |
| Ave Depth | 0.22 | 0.14 – 0.31 | <0.001 | 0.19 | 0.08 – 0.29 | <0.001 |
| **Random effects** |  |  |  |  |  |  |
| σ<sup>2</sup> | 0.20 |  |  | 0.18 |  |  |
| τ<sub>00 Lake</sub> | 0.02 |  |  | 0.03 |  |  |
| ICC | 0.10 |  |  | 0.12 |  |  |
| N<sub>Lake</sub> | 26 |  |  | 16 |  |  |
| **Other model statistics** |  |  |  |  |  |  |
| Observations | 71 |  |  | 49 |  |  |
| Mar/Con R<sup>2</sup> | 0.20 / 0.28 |  |  | 0.19 / 0.29 |  |  |

**Fig. 2.**
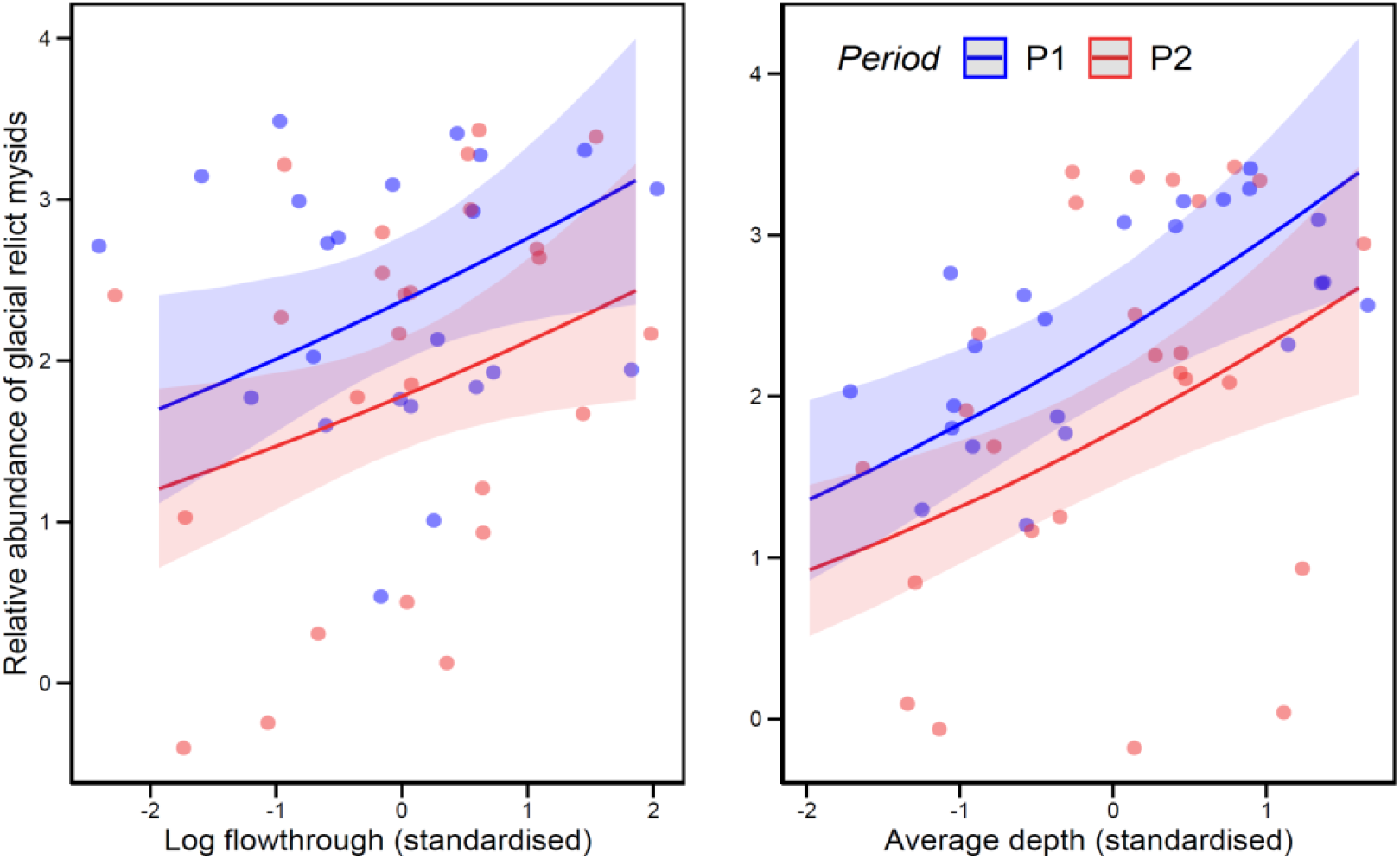
Results of the Conway-Maxwell Poisson GLMM analysis showing relationships between the relative abundance of relict mysids *Mysis relicta* against lake flowthrough and average lake depth during the 20^th^ Century (period 1, blue) and 21^st^ Century (period 2, red). Model predictions and 95% confidence intervals are shown with lines and shaded areas, raw data are shown with dots.

**Fig. 3.**
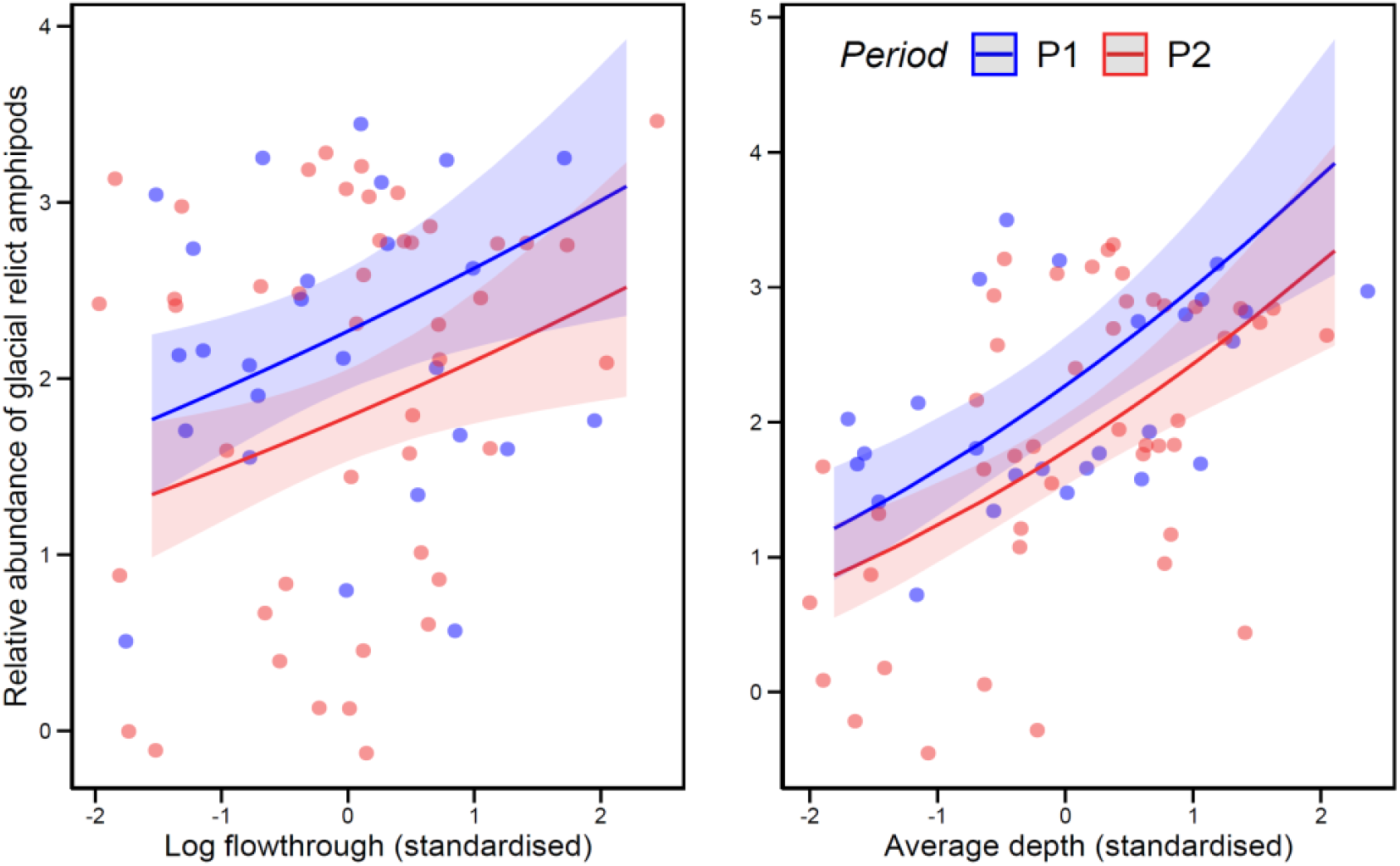
Results of the Conway-Maxwell Poisson GLMM analysis showing relationships between relative abundance of relict amphipods *Pallaseopsis quadrispinosa* against lake flowthrough and average lake depth during the 20^th^ Century (period 1, blue) and 21^st^ Century (period 2, red). Model predictions and 95% confidence intervals are shown with lines and shaded areas, raw data are shown with dots.

## 4. Discussion

Investigations of glacial relict malacostracan crustaceans in the inland waters of Lithuania started in the middle of the 20^th^ century (Gasiūnas, 1959), and historical records of their occurrence were summarised by Grigelis (1980) and Grigelis & Arbačiauskas (1996, 1997). The latest published data on the abundance and ecology of glacial relict crustaceans in Lithuania are available in Audzijonyte (1999). The study presented here provides up-to-date information on the status of these Red Data book species. Our results show a likely extinction of the glacial relict amphipod *M. affinis* in Lithuania and a rapid decrease in the number of lakes inhabited by *M. relicta* and *P. quadrispinosa*. The amphipod *M. affinis* was originally found in only two locations in Lithuania and is now also considered extinct in German and Polish waters (Köhn & Waterstraat, 1990; Żmudziński, 1995). Our data also suggests that from the 16 lakes inhabited by *M. relicta* in the 1950s, the species is now either absent or very rare in nearly half of them, i.e. in 7 lakes. Even the least sensitive species, *P. quadrispinosa*, was not found in seven lakes where it had previously been encountered.

The key contributing factor to the deteriorating conditions of glacial relict crustaceans is the eutrophication of lakes, driven mainly by agricultural run-off. Eutrophication leads to deteriorating oxygen conditions in deeper and cooler water layers, effectively reducing habitats suitable for glacial relict species. This effect is supported by our analysis, with water flowthrough and lake depth significant predictors of both relict mysid and amphipod relative abundance. Both variables are associated with the presence of cool, oxygenated water conditions. An additional possibility is that elevated water temperatures may contribute to worsening conditions for glacial relict crustaceans. However, no data are available on changes in water temperatures in the studied lakes over recent decades or how such changes might affect the summer stratification and seasonal mixing of lake water layers.

Despite initial predictions, no evidence was detected for a contribution by the presence of Ponto-Caspian crustaceans to the decreased abundance of native glacial relict crustaceans. On the one hand, this may not be surprising because glacial relicts typically occur in deep waters, while the Ponto-Caspian species occur in more shallow areas. However, if oxygen conditions in deeper waters deteriorate, relict crustaceans often occupy more shallow depths, where they may encounter competition or predation from invasive crustaceans. Despite this risk, no adverse effects of predation on glacial relicts have been seen in Lithuanian waters, and there were several lakes in which both glacial relict and Ponto-Caspian species co-existed (e.g. lakes #3, 4, 7, 16, 19, 26 for *M. relicta* in Table 1) or where glacial relicts went extinct despite there being no

Ponto-Caspian crustaceans (#5, 15, 17, 18). Similarly, predatory interactions between an invasive American amphipod *Gammarus tigrinus* and the native glacial relict mysid *Mysis salemaai* (Audzijonyte & Väinölä 2005; former *Mysis relicta*) appears not to be a threat to the native mysid in Northern Ireland (Bailey et al., 2006).

While the legacy of the Ice Age has diminished in Central Europe, Lithuanian waters still support a rich glacial faunal heritage. Nine lakes in Lithuania still harbour the relict mysid *M. relicta*, compared to just three known localities in Germany (Scharf & Koschel, 2004), up to three localities in Poland (Żmudziński, 1990) and only one in Belarus (Vezhnavets V.V., 2021, pers. comm.). This glacial heritage in Lithuania warrants preservation and improvement, where possible. The decrease in eutrophication and consequent increase in lake water quality has been shown to markedly improve environmental conditions for the relict mysids, resulting in its population increase (Scharf & Koschel, 2004).

One notable example of glacial relict extinction was in the largest Lithuanian lake, Lake Drūkšiai (#9, Table 1). Since 1984 this lake has served as a cooling reservoir for the Ignalina nuclear power plant, with the lake temperature increasing by nearly 2°C, water eutrophication level increasing due to improper wastewater discharge from the town that served the nuclear power plant, and oxygen concentrations decreasing in the lake profundal zone (Kesminas & Paškauskas, 2014; Vezhnavets & Škute, 2014). Consequently, our sampling in 2003 showed that all glacial relict species were also likely to be extinct in this lake (Table 1). However, the nuclear power plant was decommissioned in 2010, and environmental conditions in the lake are now improving, as is shown by the improved population status of two cold water species – vendace (*Coregonus albula*) and smelt (*Osmerus eperlanus*) (Kesminas & Steponėnas, 2014; Kesminas V., 2022, pers. comm.).

Given that glacial relict species cannot disperse, we would recommend attempting a re-introduction of *M. relicta* and *P. quadrispinosa* from nearby lakes, as suggested previously (Kesminas et al., 2014). A good source for translocations could be nearby Lakes Lūšiai or Aisetas, where the population status of glacial mysids and amphipods is currently good. If successful, such re-introductions could increase the probability of these endangered species surviving in Lithuania. Re-introductions into other historical habitats after an improvement in water quality should also be considered. Nevertheless, the most urgent measures to ensure the survival of the Red Data book crustaceans and other aquatic species that rely on clear and oxygenated aquatic habitats, is an urgent reduction of municipal and especially agricultural run-off into rivers and lakes.

## Acknowledgements

This study has received funding from European Regional Development Fund (Project No. 01.2.2-LMT-K-718-02-0006) under grant agreement with the Research Council of Lithuania (LMTLT).

---

**Appendix.**
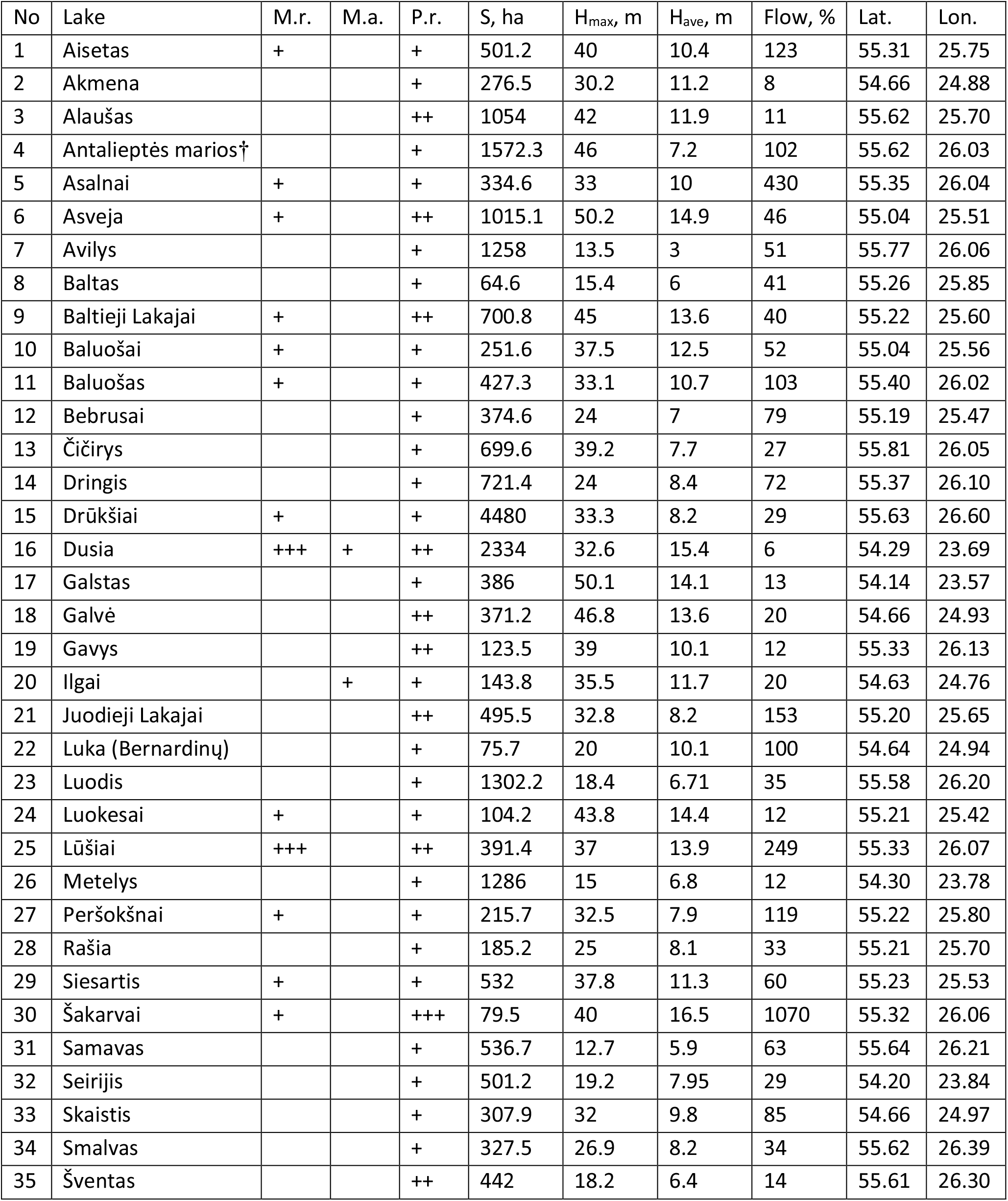
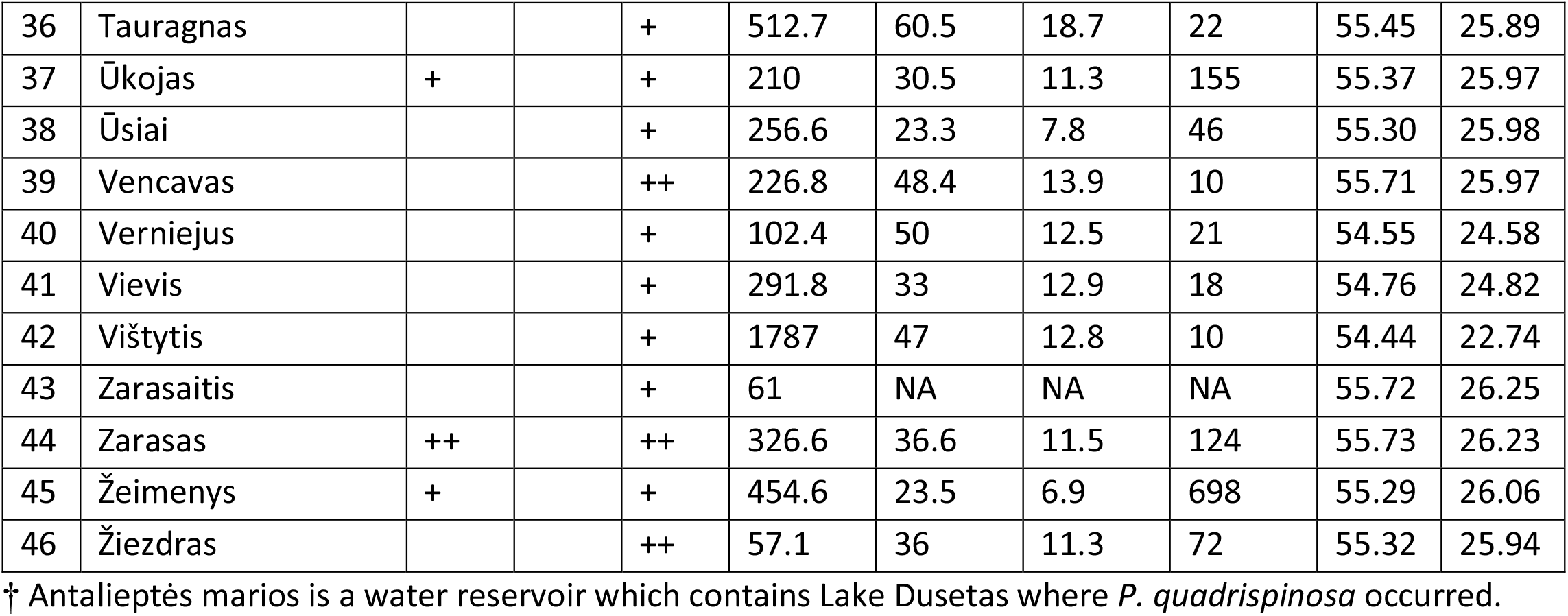
Occurrence of glacial relict malacostracan crustaceans, *Mysis relicta* (M.r.), *Monoporeia affinis* (M.a.) and *Pallaseopsis quadrispinosa* (P.q.) in Lithuanian lakes reported by Grigelis (1980) and Grigelis & Arbačiauskas (1996). Limnological characteristics: S – lake area in hectares, H_max_ and H_ave_ – maximum and average depth in meters, Flow – percentage of annual water exchange. Population status (Grigelis, 1980): +++ - abundant, ++ - moderately abundant, + - rare.

